# Comparative metagenomics reveals diversity and metabolic heterogeneity of sediment microbes from the Great Barrier Reef

**DOI:** 10.64898/2026.11.17.767448

**Authors:** Hisatake Ishida, Steven J. Robbins, Sara C. Bell, Jin Yan Lim, Marko Terzin, Nicole Webster, David G. Bourne, Patrick W. Laffy, Philip Hugenholtz, Yun Kit Yeoh, Cheong Xin Chan

## Abstract

Microbial communities in coral reefs are highly connected through benthic-pelagic coupling. Pelagic and host-associated microbiomes have recently been characterised via genome-resolved metagenomics. However, similar characterisation of sediment communities has lagged behind due to their higher complexity. Here, using short-and long-read shotgun sequence data, we characterised sediment microbial communities of Australia’s Great Barrier Reef. From 191 samples across 48 broadly distributed sites, we recovered 2,698 prokaryotic metagenome-assembled genomes (MAGs) representing 1,201 species-level clusters, most of which are novel. Compared against their sympatric counterparts in seawater, sediment MAGs exhibited greater phylogenetic diversity and flexibility in metabolic potential, implicating diverse respiration strategies and cycling of nitrogen, sulphur, and hydrogen. They also encode more abundant and novel biosynthetic capabilities, likely reflecting competition and adaptation to intermittent fluctuations in nutrient regimes, and between oxic and anoxic conditions. Our results provide foundational genomic resources for reef sediment microbes, uncovering biodiversity and function that together with planktonic microbes shape the ecological success and stability of coral reef ecosystems.

## Introduction

Coral reefs are among the most biologically diverse ecosystems on Earth. Despite high rates of primary productivity, coral reefs persist in nutrient-limited waters (de Goeij et al., 2013). This is often attributed to internal cycling of nutrients through ecological processes, such as coral-algal endosymbiosis (Muscatine & Porter, 1977), the sponge loop (de Goeij et al., 2013), and algal turf productivity to fish (Morais et al., 2020; Tebbett & Bellwood, 2021). Microbial communities are crucial to each of these processes, inhabiting diverse ecological niches across reef ecosystems, ranging from seawater to benthic hosts to sediments (Gast et al., 1999; Sorokin, 1973). These niches are highly interconnected with benthic-pelagic coupling, contributing to health and function of coral reefs (Bourne & Webster, 2013; Glasl et al., 2019). However, microbial baselines of diversity and function are still lacking for many of these niches, particularly sediments, limiting our ability to predict how reef microbial networks and the ecosystem functions they support will respond to ongoing environmental change.

Sediments are a major benthic component of coral reefs. They largely consist of calcareous sand with high permeability, porosity, and surface area (Wild et al., 2006), shaped by geochemical processes such as pore-water advection, molecular diffusion, and biophysical disturbances (Werner et al., 2006). These processes, particularly at the sediment surface, alter the availability of nutrients and oxygen over short periods and distances, creating highly dynamic microbial habitats (Gaidos et al., 2011; Hewson & Fuhrman, 2006; Uthicke & McGuire, 2007; Wild et al., 2006). Microbial communities in reef sediments process both particulate matter that settles from the water column and dissolved nutrients, subsequently exporting the resulting products to the surrounding ecosystems (Rasheed et al., 2003). These communities can include transient host-associated microbes or pathogens (Carlos et al., 2013; Nitschke et al., 2016). In these ways, sediment microbes may offer useful indicators of the health and functioning of host organisms and reef ecosystems (Glasl et al., 2019).

Metagenomic approaches are commonly used to characterise microbial diversity and functional capacity from environmental samples. Most metagenomic studies of the marine environment have focused on host-associated and pelagic microbes; few have focused on sediments of coastal and deep-sea waters (Laiolo et al., 2024). Microbial communities in marine benthic habitats, especially coral reef sediments, remain largely unexplored. Earlier research on reef sediment microbes, based on 16S rRNA gene amplicon sequencing, recovered taxa only classified at higher taxonomic ranks, belonging to phyla such as *Pseudomonadota*, *Bacteroidota*, *Cyanobacteriota*, *Planctomycetota*, *Verrucomicrobiota*, and *Acidobacteriota* (Uthicke & McGuire, 2007); taxa at lower ranks were poorly resolved using this approach, indicating potential microbial novelty in these systems. In comparison, shotgun metagenomics can characterise microbial diversity at greater taxonomic resolution (Brumfield et al., 2020). Metagenomic data also reveal metabolic potential through gene-centric analysis of individual gene sequences or genome-centric analysis of metagenome-assembled genomes (MAGs; Hugenholtz & Tyson, 2008). Although such analyses of soil/sediment microbes are known to be complicated by high biomass, taxonomic diversity, and spatial heterogeneity (Nayfach et al., 2021; van der Walt et al., 2017), the use of long-read sequencing is proving to better capture this inherent complexity (Sereika et al., 2025).

Here, adopting a shotgun metagenomic approach incorporating both short-and long-read sequencing technologies, we assessed the diversity and metabolic potential of prokaryotic communities in reef sediments from 48 sites spanning seven geographic sectors across Australia’s Great Barrier Reef (GBR). By comparing these data against those generated from seawater collected at the same sites and times (Robbins et al., 2026; Terzin et al., 2025; Terzin et al., 2026), we reveal niche-specific communities and functional capacity, with greater phylogenetic and metabolic diversity observed among the sediment microbes. This study expands the current GBR Microbial Genomes Database (GBR-MGD), which has focused on seawater datasets (hereinafter, “MGD-seawater”; Robbins et al., 2026), through the addition of reef sediment-derived genomic resources (“MGD-sediment”). Together, these complementary resources provide a foundational reference for investigating contributions of free-living microbial communities to the function and resilience of the GBR, and, more broadly, global coral reef ecosystems.

## Materials and Methods

### Sample collection and processing

Surface sediments were collected from 48 sites (in four replicates) across four sampling trips between November 2019 and July 2020 (total 192 samples; Table S1). Spanning the length of the GBR (Fig. 1a), sampling was conducted with the Australian Institute of Marine Science’s Long-Term Monitoring Program surveys. The first three trips occurred during the austral summer, covering 34 sites: the northern GBR (Trip 1 in November-December 2019; Cape Grenville and Princess Charlotte Bay sectors), the southern GBR (Trip 2 in January 2020; Swain and Capricorn Bunker sectors), and the central GBR (Trip 3 in February 2020; Cairns and Innisfail sectors). The remaining 14 sites in the central GBR were sampled during the austral winter (Trip 4 in July 2020; Townsville sector). These 48 sites were selected to include reefs of contrasting zoning status within the GBR Marine Park: 23 sites from no-take marine reserves (all extractive activities are prohibited) and 25 from reefs open to fishing (fished reefs: 5 yellow and 20 dark blue zones; Terzin et al., 2026). Samples were processed in 2 × 1 g replicates for “shallow” Illumina sequencing (165 samples), or 6 × 1 g replicates for “deep” Illumina and nanopore sequencing (27 samples). For each sample, microbial cells were detached from sediment particles using a cell separation buffer (0.35% polyvinylpyrrolidone (w/v), 0.13% sodium pyrophosphate (w/v), and 0.5% Tween 20 (v/v) in calcium-and magnesium-free seawater), followed by sonication (50 s, 50% amplitude) and low-speed centrifugation (300 × *g*, 5 min). The resulting supernatants were prefiltered through 8 µm and 5 µm filters in Swinnex housings (Millipore) to remove large debris. Microbial cells were then collected on 0.22-µm Sterivex-GP filters (Millipore).

**Fig. 1.**
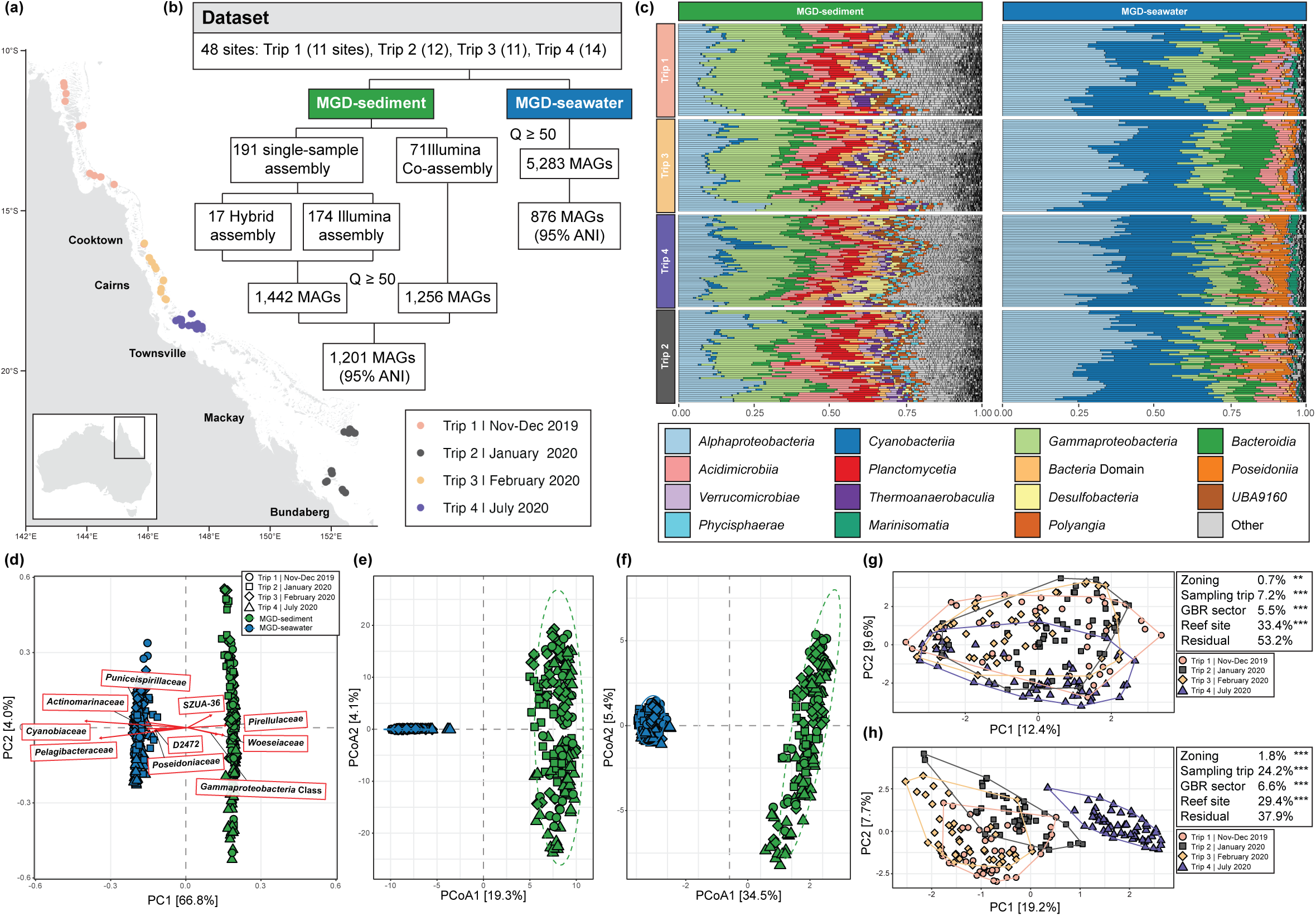
Overview of metagenomic datasets generated from sediment and seawater microbes of the Great Barrier Reef. (a) Map of the Great Barrier Reef showing the locations of 48 sampling sites. Points represent each site, and are coloured by sampling trip. (b) Summary of metagenomic datasets generated from MGD-sediment (green) and MGD-seawater (cyan). Only metagenome-assembled genomes (MAGs) with quality score Q (genome completeness − 3 × contamination) ≥ 50% were retained. (c) Relative abundance of the top 15 prokaryotic classes in sediment and seawater communities, identified based on operational taxonomic units derived from phylogenetic marker genes (mOTUs). Samples (rows) are grouped by habitat and sampling trip. (d) Principal Component Analysis (PCA) of mOTU-based prokaryotic community profiles at the family level for sediment and seawater communities. Points represent each sample, and are shaped by sampling trip. Only the ten taxa with the longest loading vectors are shown. (e-f) Principal Coordinate Analysis (PCoA) of normalised reads per kilobase per million (RPKM) abundance of KO-annotated (e) and CAZyme-annotated (f) genes in sediment and seawater communities. (g-h) PCA of mOTU-based prokaryotic community profiles at the family level for sediment (g) and seawater (h) communities. The accompanying table shows the proportion of variation in community composition explained by reef zoning status, sampling trip, GBR sector, and reef site, as inferred by sequential PERMANOVA (***: *p* < 0.001; **: 0.001 ≤ *p* < 0.01).

DNA was extracted from the 0.22-µm Sterivex filters following a protocol modified from Robbins et al. (2026). Each filter was filled with 1.8 mL of filter-sterilised lysis buffer containing 50 mM Tris-HCl (pH 8.0), 40 mM EDTA (pH 8.0), 256 mg·mL^−1^ sucrose and 36 µL lysozyme (100 mg·mL^−1^). The filters were incubated with mild agitation (1 h, 37°C), followed by the addition of 40 µL proteinase K (10 mg·mL^−1^ in SDS) and further incubation (1.5 h, 65°C). The lysate was then ejected through the Sterivex filter into a 5-mL microtube, and proteins were precipitated by adding one-third volume of 5 M potassium acetate, followed by incubation on ice for 30 min and centrifugation (14,000 × *g*, 15 min). The supernatant was transferred to a new tube, and DNA was precipitated with 0.8 volumes of isopropanol for 15 min at room temperature and was recovered by centrifugation (20,000 × *g*, 30 min, 4°C). The pellet was resuspended in 400 µL TE buffer, and the DNA was purified with an equal volume of chloroform:isoamyl alcohol (24:1 v/v) and centrifugation at 16,000 × *g* for 10 min. DNA in the aqueous phase was precipitated again with 0.8 volumes of isopropanol and recovered by centrifugation (20,000 × *g*, 25 min, 4°C). The DNA was washed twice with 70% ethanol, dried in a vacuum concentrator, resuspended in 30 µL TE buffer overnight at 4°C, and stored at −75°C.

### Shotgun metagenomic sequencing

For each sample, a sequencing library was prepared using Nextera DNA Flex Library Preparation Kit (Illumina #20018705) and pooled at equimolar amounts of 2 nM. For “shallow” Illumina sequencing at a target depth of 5 Gbp, 164 libraries were sequenced (2 × 150 bp paired-end) on the Illumina NovaSeq 6000 platform with S4 flow cells at the Centre for Microbiome Research (Brisbane, Australia) and Microba Life Sciences (Brisbane, Australia). For 27 sites, one replicate was selected for “deep” Illumina sequencing at a target depth of 40 Gbp, and was also processed for nanopore long-read sequencing; both were conducted at the Australian Centre for Ecogenomics sequencing facility (Brisbane, Australia). For the long-read sequencing, short DNA fragments (< 10 kb) were first removed using Circulomics SRE XS kit (PacBio, SKU 102-208-200), and libraries were prepared from the size-selected DNA using Ligation Sequencing Kit (Oxford Nanopore, LSK-109) with PCR-Barcoding Expansion Kit (EXP-PBC001). Barcoded libraries were pooled (two samples per flow cell at equal concentration) and sequenced on Oxford Nanopore Technologies (ONT) PromethION 24 using R9.4 flow cells for 72 hours with MinKNOW v20.06.18 under default settings. Guppy v5.0.16 was then used with the super-accuracy model to perform basecalling.

### Metagenomic assembly, binning and quality control

In total, we generated Illumina short-read sequencing data from 191 samples (DNA from one sample from reef 11-162 did not meet requirements for sequencing). While all 27 “deep” samples yielded high-quality Illumina short-read data (mean yield of 255 million paired-end reads and 96% of bases with Q ≥ 30), only 17 yielded ONT data of sufficient quality with a mean yield of 9 million reads (8 Gbp) per sample. Sequencing adapters were removed using fastp v0.23.4 (Chen et al., 2018) and porechop v0.2.4 (Wick, 2017) for Illumina and ONT datasets, respectively.

For the 17 samples for which both Illumina and acceptable long-read data were available, assemblies were generated using Aviary *assemble* v0.9.1 (Newell et al., 2024), with metaFlye v2.9.4 (Kolmogorov et al., 2020) for ONT reads, and metaSPAdes v3.15.4 (Nurk et al., 2017) for Illumina reads. metaSPAdes v3.15.4 was used to assemble all the other 174 samples for which only Illumina data were available. In total, 191 single-sample assemblies were generated (Fig. 1b). For each assembly, 20 Illumina datasets were selected for multi-sample binning based on similarity in microbial diversity between the target assembled contigs and Illumina reads (Table S2). This was calculated based on a profile of 59 single-copy marker genes (22 bacteria-specific, 24 archaea-specific, and 13 common to both domains) recovered using *pipe* and *appraise* of SingleM v1.18.0 (Woodcroft et al., 2025), with a reference metapackage built with Genome Taxonomy Database (GTDB) Release 09-RS220 (Parks et al., 2022). Multi-sample binning was performed using Aviary *recover* v0.9.1. Briefly, the *recover* pipeline employs DAS Tool v1.1.2 (Sieber et al., 2018) to select the best representative MAGs produced by multiple binning algorithms, including MetaBAT1 v2.15 (Kang et al., 2015), MetaBAT2 v2.15 (Kang et al., 2019), SemiBin v2.1.0 (Pan et al., 2022), VAMB v3.0.2 (Nissen et al., 2021), and Rosella v0.5.2 (Newell et al., 2023).

To maximise recovery of MAGs from the 191 Illumina datasets, we also adopted a co-assembly strategy. Pairs of samples with similar microbial composition were first identified using BinChicken v0.12.11 *coassemble* (Aroney et al., 2025a) based on profiles of the 59 marker genes as above, and co-assembled using MEGAHIT v1.2.9 (Li et al., 2015) implemented within the Aviary *assemble* pipeline. In total, 71 co-assemblies were generated (Table S2; Fig. 1b). For each co-assembly, 20 Illumina datasets were selected for co-binning by BinChicken *coassemble* (Table S2).

The quality of reconstructed MAGs was evaluated using both CheckM v1.2.2 (Parks et al., 2015) and CheckM2 v1.0.2 (Chklovski et al., 2023); MAGs were retained for downstream analyses if either tool reported a quality score (i.e. genome completeness minus three times contamination) of ≥ 50%, following Robbins et al. (2026). In total, 2,698 MAGs were recovered from these approaches, and further dereplicated at 95% average nucleotide identity (ANI) using CoverM *cluster* v0.7.0 (Aroney et al., 2025b) with the ‘Parks2020_reduced’ formula, providing approximate species-level clustering. This step yielded 1,201 “species-resolved” MAGs overall from the sediments.

### Genome-centric analyses for assessing taxonomic and metabolic diversity

These 1,201 sediment MAGs, alongside 876 seawater MAGs (MGD-seawater datasets; https://zenodo.org/records/20503966) recovered from the co-located surface seawater (Robbins et al., 2026), were taxonomically annotated using *classify_wf* of GTDB-Tk v2.4.1 (Chaumeil et al., 2019) based on GTDB Release 10-RS226 (Parks et al., 2026). Following methods implemented in GTDB-Tk (Chaumeil et al., 2019), assigned taxonomy was curated through manual inspection of the genome trees generated using *de_novo_wf* of GTDB-Tk, followed by a comparison of relative evolutionary distance (RED) assigned at each node with median RED values calculated for each taxonomic rank using PhyloRank v0.1.12 (https://github.com/dparks1134/PhyloRank). To assess diversity of the combined 2,077 MAGs, a phylogeny was inferred from phylogenetically informative marker genes, independently for bacteria based on the bac120 marker-gene set of GTDB-Tk, and for archaea based on the arc53 set. Multiple sequence alignment of concatenated marker genes was used to infer a maximum likelihood tree using IQ-TREE v2.2.2.3 (Minh et al., 2020) with 1,000 ultrafast bootstrap replicates, specifying LG + G model for the bacterial tree, and LG + C10 + F + G model for the archaeal tree. To assess taxonomic novelty, sediment and seawater MAGs were compared against a database of publicly available genomes using CoverM *cluster* with 95% ANI and the ‘Parks2020_reduced’ formula. This database encompasses prokaryotic MAGs from publicly available data repositories: the Ocean Microbiomics Database (OMD; Paoli et al., 2022), Genomes from Earth’s Microbiomes (GEMs; Nayfach et al., 2021), shallow-water coastal marine sediments (Silva-Solar et al., 2024; Viver et al., 2025), and reef sediments from the South China Sea (Dong et al., 2023). Community profiling and genome recovery were assessed by mapping short-read datasets from each habitat separately to the corresponding MAGs (sediment reads to 1,201 sediment MAGs and seawater reads to 876 seawater MAGs) using CoverM *genome* with exclusion of read alignments with < 95% identity and < 75% alignment length.

Sediment and seawater MAGs were functionally annotated using DRAM v1.5 (Shaffer et al., 2020) with KOfam (Aramaki et al., 2020), Pfam (Mistry et al., 2021), and MEROPS (Rawlings et al., 2018) databases. CAZyme families and CAZyme Gene Clusters (CGCs) were identified using run_dbcan v5.0.1 (Zheng et al., 2023), with substrate target of CGCs inferred based on their similarity with polysaccharide utilisation loci (PULs) in dbCAN-PUL database (Ausland et al., 2021). Further annotations were performed through homology-based searches using DIAMOND blastp v2.1.0 (Buchfink et al., 2021) against SulfAtlas database v2.3.1 (Stam et al., 2023) for sulphatases, HydDB (Søndergaard et al., 2016) for 30 hydrogenase subgroups, and a custom protein database of 44 metabolic marker genes (Greening et al., 2019; Leung et al., 2026; Ortiz et al., 2021), containing key enzymes for aerobic and anaerobic respiration, energy conservation from organic/inorganic compounds, nitrogen cycling, sulphur cycling, carbon cycling, and phototrophy. GraftM v0.15.1 (Boyd et al., 2018) was also used to identify ammonia monooxygenase (*amoA*; GraftM package 7.22), and to classify these sequences into functional clades (i.e. bacterial or archaeal *amoA*). To examine gene-copy enrichment, annotations of 796 highly complete MAGs (completeness ≥ 85%; 347 from the sediment, 449 from seawater) were compared using Mann-Whitney U-test in the stats v4.3.1 R package for KOs, CAZymes, Pfams, MEROPS, SulfAtlas, HydDB, and a custom list of 44 metabolic marker genes, followed by Benjamini-Hochberg multiple-testing correction (Benjamini & Hochberg, 1995).

Biosynthetic gene clusters (BGCs) were identified using antiSMASH v6.1.1 (Blin et al., 2021) with a minimum contig length of 5 kbp for 1,201 sediment MAGs, 876 seawater MAGs (Robbins et al., 2026), 8,422 MAGs (dereplicated at 95% ANI from 34,815 MAGs) of OMD (Paoli et al., 2022), and 3,899 MAGs (dereplicated at 95% ANI from 13,549 MAGs) of the Reef Microbiomics Database (RMD; Wiederkehr et al., 2026). Alongside the experimentally validated BGCs from MIBiG v4.0 (Zdouc et al., 2025), these predicted BGCs were clustered into gene cluster families (GCFs) using clust-o-matic (Gavriilidou et al., 2022) at a clustering threshold of 0.5. antiSMASH annotations of the individual BGCs were parsed to assign a natural product class for each GCF, as defined in BiG-SCAPE (Navarro-Muñoz et al., 2020). The novelty of GCFs was estimated based on L2-normalised distances to BGCs within BiG-FAM (Draisma et al., 2026; Kautsar et al., 2021), consisting of computationally predicted BGCs from ∼190,000 genomes deposited in RefSeq (O’Leary et al., 2016). A GCF was identified as novel if the mean distance (*d*) of its member BGCs to their closest GCFs in BiG-FAM was above 0.4.

### Gene-centric analyses for assessing taxonomic and metabolic diversity

Illumina reads from sediment and seawater datasets were used for gene-centric analysis of taxonomic diversity. To account for uneven sequencing depth, all datasets were subsampled to 10 million paired-end reads. SingleM *pipe* was used to generate prokaryotic community profiles based on the recovery of operational taxonomic units derived from phylogenetic marker genes (mOTUs). Principal Component Analysis (PCA) was applied to the centred log-ratio (CLR)-transformed profiles using the microViz v0.12.7 R package (Barnett et al., 2021) to visualise major compositional differences between sediment and seawater communities. PCA was subsequently performed separately for sediment and seawater datasets to assess within-habitat community structure. SingleM *pipe* was also run on assembled contigs, reconstructed MAGs, and 143,614 representative genomes of GTDB Release 10-RS226. SingleM *appraise* was then used to trace the loss of community representation during assembly and binning steps, and to assess the extent of marker gene novelty compared to GTDB Release 10-RS226. SingleM *microbial_fraction* was used to predict the prokaryotic proportion of the sequenced microbial community. Assembled contigs were also taxonomically annotated using MMseqs2 v15.6f452 taxonomy module (Steinegger & Söding, 2017) against a custom database comprising protein sequences deposited in RefSeq, EukProt (Richter et al., 2022), PLSDB (Galata et al., 2019), IMG/VR (Paez-Espino et al., 2017), as well as those predicted from genomes in GTDB Release 08-RS214 (Robbins et al., 2026).

To assess metabolic functions of the microbial community, a non-redundant gene catalogue was generated by searching for complete open reading frames (ORFs) from all assembled contigs using Prodigal v2.6.3 (Hyatt et al., 2010), and clustering them at > 90% identity across > 80% of protein length using MMseqs2 *easy-linclust* module. In total, 60,940,012 complete ORFs were clustered into 39,225,676 non-redundant protein sequences, which were functionally annotated using DRAM *annotate_genes* for KOfam genes, run_dbcan *CAZyme_annotation* for CAZyme families, and GraftM (package 7.22) for *amoA*. DIAMOND blastp searches were also performed against HydDB and the same curated 44 metabolic marker gene sets used for genome annotation. Rarefied Illumina reads were aligned to these sequences using DIAMOND blastx to estimate protein abundance, applying filters for read length (≥ 140 bp) and alignment parameters (*--evalue 0.00001 --min-score 40 --query-cover 80 --id 70 --max-hsps 1 --max-target-seqs 1*), and removing low-abundance proteins (represented by < 0.0001% reads). Reads were also aligned to the single-copy marker genes included within the SingleM GTDB Release 10-RS226 metapackages using the same settings. The resulting read counts were normalised to reads per kilobase per million (RPKM) values for both the protein database and SingleM marker genes. RPKM values of the protein database were then normalised based on the mean RPKM value of all SingleM marker genes on a per-sample basis for comparability. The resulting normalised RPKM matrix was transformed into an Aitchison distance matrix and visualised by Principal Coordinate Analysis (PCoA) using microViz. Differential abundance analysis with reef site as a random effect was conducted on mOTU profiles and functional profiles (based on KOs, CAZymes, hydrogenases, and above-described custom 44 marker gene sets) using MaAsLin2 v1.16.0 (Mallick et al., 2021) and LinDA v0.2.0 (Zhou et al., 2022); results were considered significant if supported by both methods.

### Community structure and reef zoning status analyses

Community structure was assessed using mOTU-based profiles (aggregated at the family level) of both habitats and MAG-based profiles of sediment communities; the corresponding MAG-based analyses of seawater communities were reported by Terzin et al. (2026). Sequential PERMANOVA was performed on Aitchison distances calculated from CLR-transformed abundance data with 9,999 permutations for significance testing (model formula: Aitchison distances ∼ reef zoning status + sampling trip + GBR sector + reef site). To test whether reef zoning status could be predicted from sediment community composition, we followed the approach of Terzin et al. (2026), using Multivariate INTegration sparse Partial Least Squares Discriminant Analysis (MINT sPLS-DA) in mixOmics v6.26.0 (Rohart et al., 2017), with GBR sector as the study variable to account for sector-specific variation. The number of features retained per component was tuned using Leave-One-Group-Out Cross-Validation (LOGOCV; test values of 10 to 500 in steps of 10; balanced error rate; centroids distance), and the final two-component model was evaluated using the same cross-validation scheme. Classification performance was assessed using balanced accuracy and cross-validated area under the receiver operating characteristic curve. The direction of enrichment for each selected feature was determined from the difference in mean CLR abundance between no-take marine reserves and fished reefs. To assess statistical significance, zoning status labels were permuted 999 times among reefs within each GBR sector, and the MINT sPLS-DA model was refitted with the tuned parameters and evaluated by LOGOCV to generate a null distribution of classification accuracy.

## Results and Discussion

### Microbial communities in sediments are more diverse, complex, and novel than those in surrounding seawaters

We generated shotgun metagenomic sequence data from microbial communities (0.22-5 µm size fraction) of 191 coral reef sediments, collected from 48 sites across the GBR (Materials and Methods; Fig. 1a-b), constituting the MGD-sediment datasets. The mean yield per sample for Illumina short-read sequencing was 21 Gbp (total 4 Tbp); ONT long-read data were generated from a subset of 17 samples, with a mean yield of 8 Gbp (mean read length 911 bp; total 130 Gbp; Table S3). These complement the MGD-seawater datasets generated from surface seawaters collected at the same times and sites, representing the sympatric planktonic microbial communities (Laffy et al., 2022; Robbins et al., 2026; Terzin et al., 2025). Hereinafter, we consider sediments and surface seawaters as two distinct microbial habitats.

Based on analysis of mOTUs, we detected 2,296 prokaryotic taxa (2,172 Bacteria, 124 Archaea) across all samples, with Bacteria dominating the prokaryotic fraction of the community (mean relative abundance 97.4 ± 1.3% in sediments, 95.5 ± 3.7% in seawaters). At the class level (Fig. 1c), the most abundant taxa in sediments were from *Gammaproteobacteria* (20.6 ± 3.3%), followed by *Alphaproteobacteria* (13.3 ± 5.4%), *Acidimicrobiia* (8.3 ± 4.0%), *Planctomycetia* (7.0 ± 2.2%), and *Bacteroidia* (6.0 ± 2.4%). By contrast, the top-ranking classes in seawater samples were *Alphaproteobacteria* (35.6 ± 7.8%), *Cyanobacteriia* (28.5 ± 10.1%), *Bacteroidia* (9.6 ± 4.4%), *Gammaproteobacteria* (9.6 ± 2.8%), and *Acidimicrobiia* (5.9 ± 2.4%). This compositional difference was also observed in the PCA ordination of community profiles (aggregated at the family level where possible), in which sediment and seawater communities formed distinct clusters along PC1 (Fig. 1d). This habitat-specific pattern was associated with the increased relative abundances of the families *Pirellulaceae* (*Planctomycetia*; mean 4.37% relative abundance in sediment versus 0.04% in seawater), *Woeseiaceae* (*Gammaproteobacteria*; 3.30% versus 0.01%), and *SZUA-36* (*Gammaproteobacteria*; 1.93% versus 0.00%) in sediments (Tables S4 to S6), whereas seawater communities were dominated by the bacterioplankton families *Cyanobiaceae* (*Cyanobacteriia*; 28.18% versus 0.11% in sediment) and *Pelagibacteraceae* (*Alphaproteobacteria*; 19.19% versus 0.00%). These families were also highly prevalent across their respective habitat types (> 80%; Tables S4 to S6), whereas *Flavobacteriaceae*, *Rhodobacteraceae*, and *Halieaceae* were prevalent in both habitats (> 80% of all samples). Alpha-diversity at family level was significantly higher in sediment communities compared to those of seawaters (mean Shannon index = 4.47 ± 0.23 versus 2.74 ± 0.25; *p* < 2 × 10⁻^16^ in Wilcoxon rank-sum tests). Dissimilarity between the communities was also evident in the functional profiles of KO-annotated (Fig. 1e) and CAZyme-annotated (Fig. 1f) genes.

Community composition in each habitat revealed distinct clustering patterns when assessed independently based on PCA ordination. Sediment communities showed no clear clustering by temporal factors (sampling trips or season) or by spatial gradients (GBR sectors) captured by our sampling design (Fig. 1g). In contrast, seawater communities were primarily clustered by season (Fig. 1h), with samples collected during austral winter (Trip 4) separated from those collected during austral summer (Trips 1 to 3); this follows earlier observations, where seawater communities differed between summer and winter but not among summer trips spanning the latitudinal extent of the GBR (Terzin et al., 2025). These patterns were largely supported by sequential PERMANOVA (Fig. 1g-h). In both habitats, reef site explained the largest proportion of community variation (sediment 33.4%, seawater 29.4%; both *p* < 0.001) after accounting for zoning status, sampling trip and GBR sector. The two habitats differed in the effect of sampling trip, which explained 24.2% of the variation in seawater communities but only 7.2% in sediment communities. This exceeded the effects of GBR sector (seawater 6.6%, sediment 5.5%) and reef zoning status (seawater 1.8%; sediment 0.7%). A larger proportion of variation remained unexplained in sediment than in seawater communities (seawater 37.9%, sediment 53.2%). Previous studies have shown that reef sediment microbes are highly site-specific and structured by local habitat characteristics (e.g. sediment grain size and chemical composition; Glasl et al., 2019; Olmos Pin et al., 2026). Although these were not measured here, the dominant effect of reef site and the large unexplained variation in sediment communities likely reflect such fine-scale, reef-specific drivers. Because each reef was sampled once and winter samples were collected from a single region (Townsville sector), sampling trip captures both seasonal and regional variation, and our snapshot design cannot strictly separate the two. Nevertheless, sediment communities showed weaker differentiation among trips than seawater communities, indicating lower sensitivity to broader seasonal and regional variation.

These patterns nonetheless may need to be interpreted with caution due to the underrepresentation of reef sediment taxa in current reference databases. When compared against mOTUs derived from all genomes in the GTDB Release 10-RS226 database, only 1.4% of bacterial and 0.1% of archaeal sediment mOTUs could be clustered at species level, compared to 35.7% and 26.7% respectively for the seawater mOTUs. This disparity indicates greater taxonomic novelty among sediment communities relative to co-located reef seawater communities, and to other ecosystems currently represented in GTDB. These observations highlight the importance of reconstructing microbial genomes from reef sediments, not only for resolving the ecology and metabolic potential of individual taxa, but also for expanding the ecological and taxonomic breadth of reference databases used in microbial community analyses.

### MAGs recovered from sediments capture novel and abundant taxa

From the sediment datasets, 191 single-sample assemblies (17 hybrid and 174 Illumina-only assemblies) and 71 co-assemblies were generated (Table S2; Fig. 1b); from these assemblies, 2,698 prokaryotic MAGs were recovered and dereplicated, yielding 1,201 bacterial and archaeal genomospecies (Figs 2a and S1; Materials and Methods), with mean completeness of 74.17 ± 13.35% and contamination of 2.42 ± 2.00% (CheckM2; Tables S7 and S8). Almost all genomospecies (1,199) are bacterial, collectively belonging to 27 phyla (Fig. 2b); the most diverse are *Pseudomonadota* (331 *Gammaproteobacteria* and 201 *Alphaproteobacteria*), *Actinomycetota* (266), *Acidobacteriota* (82), *Bacteroidota* (61), *Desulfobacterota* (34), and *Planctomycetota* (34). The two archaeal genomospecies belong to *Thermoproteota*. This observation aligns with earlier descriptions of community compositions in GBR sediments based on 16S rRNA gene sequencing (Uthicke & McGuire, 2007). However, 1,182 (98% of 1,201) genomospecies we recovered do not have representative genomes in GTDB Release 10-RS226 (Fig. 2c; Table 1), and 1,158 (96% of 1,201) genomospecies could not be clustered with other publicly available genomes (Materials and Methods) at species level (Fig. 2d; 95% ANI). These MAGs may therefore represent either newly recognised reef-associated taxa, potentially specific to GBR sediments, or taxa that are present elsewhere but remain underrepresented in public databases due to difficulties in genome reconstruction.

**Fig. 2.**
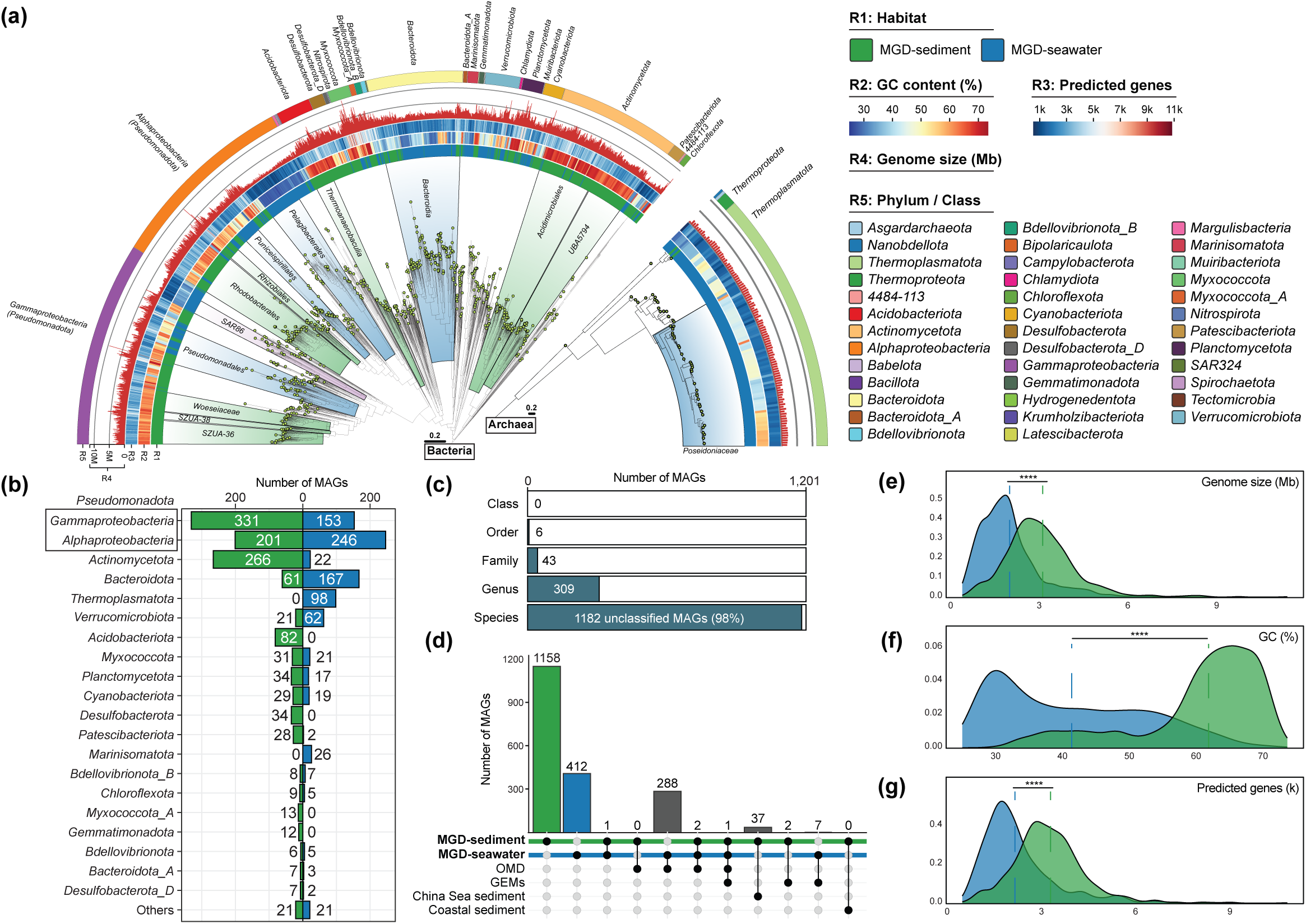
**Genomic features of metagenome-assembled genomes (MAGs) from sediment and seawater microbes of the Great Barrier Reef. (**a) Maximum-likelihood phylogeny of bacterial and archaeal MAGs with quality score (completeness − 3 × contamination) ≥ 50%, recovered from sediment (1,201 MAGs) and seawater (876 MAGs) samples of the GBR. MAGs with completeness ≥ 85% are indicated by yellow circles at branch tips. Clades preferentially represented by a greater number of MAGs in either habitat are highlighted from their last common ancestor (green for sediment and cyan for seawater). Rings indicate, from inner to outer: MAG source (R1; sediment or seawater), GC content (R2; as percentage), number of predicted genes (R3), genome size (R4; in Mb), and phylum-level taxonomic affiliation (R5; *Pseudomonadota* split into *Alphaproteobacteria* and *Gammaproteobacteria*). (b) Number of MAGs assigned to each phylum (or class for *Pseudomonadota*), recovered from sediment (left) and seawater (right). (c) Taxonomic novelty of sediment MAGs across taxonomic ranks (from class to species) according to Genome Taxonomy Database Release 10-RS226; classified fraction in white and unclassified in blue. (d) Species-level comparison (95% ANI) of sediment MAGs against public genome collections, including MGD-seawater, Ocean Microbiomics Database (OMD), Genomes from Earth’s Microbiomes (GEMs), South China Sea reef sediment, and Isfjorden and Helgoland coastal sediment. Distribution of genome size in Mb (e), GC content (f), and number of predicted genes (g) for sediment and seawater MAGs. Dashed lines indicate mean values for each genomic feature, coloured by habitat (****: *p* ≤ 0.0001).

**Table 1.** Taxonomic novelty of metagenome-assembled genomes retrieved from sediment and seawater samples across the Great Barrier Reef.

|  | MGD-sediment |  | MGD-seawater |  |
|---|---|---|---|---|
| Taxonomic rank | # of taxa | # of novel taxa | # of taxa | # of novel taxa |
|---|---|---|---|---|
| Domain | 2 | 0 | 2 | 0 |
| Phylum | 28 | 0 | 25 | 0 |
| Class | 56 | 0 | 38 | 0 |
| Order | 120 | 4 (3.3%) | 84 | 0 |
| Family | 220 | 25 (11.4%) | 145 | 6 (4.1%) |
| Genus | 539 | 221 (41.0%) | 326 | 42 (12.9%) |
| Species | 1,201 | 1,182 (98.4%) | 876 | 517 (59.0%) |

Mapping of short-read data to the 1,201 sediment MAGs recruited only a small proportion of reads (0.48−18.35% of reads per sample; mean 4.43%); > 95% of reads did not map to any MAGs in 132 (69% of 191) samples. Taxonomic annotation of unbinned contigs showed a dominance of prokaryotic sequences (mean 91.54% from Bacteria, 0.53% from Archaea), with limited representation of non-prokaryotic sources, i.e. of viral (1.06%) and eukaryotic (0.44%) origins. These results highlight the challenges in reconstructing genomes from highly diverse communities comprising abundant closely related and/or heterogeneous strains, an issue known for metagenomic analysis of soil and sediment samples (Howe et al., 2014). To further evaluate the extent to which the recovered MAGs represent the prokaryotic communities in our sediment samples, we assessed community profiles using a genome-centric approach based on MAGs, and a gene-centric approach based on recovery of single-copy marker genes (Materials and Methods). Both approaches revealed *Gammaproteobacteria*, *Alphaproteobacteria*, *Acidimicrobiia*, and *Bacteroidia* as the dominant classes based on relative abundances (Fig. S2; Table S9), suggesting that the recovered MAGs represent some of the abundant prokaryotic taxa in the sediments. However, some taxa including the family *Pirellulaceae* (*Planctomycetia*) were underrepresented in the genome-centric profiles, possibly reflecting the difficulty of recovering their large and complex genomes (Lage et al., 2025).

### Sediment and seawater MAGs exhibit habitat-specific diversity, distinct genome features, and differing sensitivity to reef zoning

To assess diversity of MAGs reconstructed from both habitats (Table S8), we inferred maximum likelihood trees for bacterial and archaeal taxa, based on marker genes recovered in the genomes (see Materials and Methods). This resulted in a 1,972-taxon bacterial tree and a 105-taxon archaeal tree, revealing distinct microbial diversity in sediment versus seawater (Fig. 2a). Only four bacterial taxa (*RCC307 sp012270465*, *UBA1268 sp002694955*, *Parasynechococcus sp002724845*, and *Hepatobacter* sp.) were recovered from both habitats. Consistent with the gene-centric mOTU profiles, nearly half of the MAGs represent *Pseudomonadota* (Figs 2a-b and S1). Among *Gammaproteobacteria*, the orders *Xanthomonadales* (74 MAGs from *SZUA-36* and 24 from *SZUA-38*) and *Woeseiales* (all 60 MAGs from *Woeseiaceae*) were prevalent and found only in the sediments, whereas *Pseudomonadales* (66 MAGs) and *SAR86* (45 MAGs) were common in the seawaters (Fig. 2a). Among *Alphaproteobacteria*, the orders *Rhodobacterales* (all 111 MAGs from *Rhodobacteraceae*) and *Rhizobiales* (33 MAGs) were common in the sediments, whereas *Pelagibacterales* (99 MAGs; 97 from *Pelagibacteraceae*) and *Puniceispirillales* (52 MAGs) were prevalent and found only in the seawaters (Fig. 2a). The second most represented phylum among the sediment MAGs was *Actinomycetota* (Fig. 2b), particularly within orders *Acidimicrobiales* (151 MAGs) and *UBA5794* (59 MAGs); *UBA5794* was detected only in sediments. This compares to only 22 *Actinomycetota* MAGs recovered from the seawater. The second most represented phylum among the seawater MAGs was *Bacteroidota* (Fig. 2b), all represented by class *Bacteroidia*. Other habitat-specific taxa included *Acidobacteriota* and *Desulfobacterota* in the sediments, versus *Marinisomatia* and the archaeal phylum *Thermoplasmatota* in seawaters.

These taxonomic differences were accompanied by distinct genome features. Compared with seawater MAGs, sediment MAGs exhibited significantly larger genomes (mean 3.1 Mb versus 2.0 Mb; Fig. 2e), higher GC content (mean 60.6% versus 41.5%; Fig. 2f), and more predicted genes per genome (mean 3,252 versus 2,284; Fig. 2g); all comparisons yielded *p* < 2 × 10⁻^16^ in Wilcoxon rank-sum tests. These differences in genomic properties are consistent with contrasting nutrient regimes between the two habitats (Lauro et al., 2009). In relatively copiotrophic habitats such as sediments, microbes may benefit from larger genomes, harbouring more diverse gene repertoires that enable the use of a wide range of available resources (Bentkowski et al., 2015). Genome expansion has previously been associated with increased metabolic versatility and pangenome diversification in terrestrial soil microbes (Maistrenko et al., 2020; Nayfach et al., 2021). In contrast, microbes inhabiting oligotrophic environments such as seawater may benefit from smaller and more streamlined genomes, reducing the energetic and nutrient costs of genome replication and cellular maintenance (Giovannoni et al., 2014). Members of the *Pelagibacteraceae*, which were prevalent and abundant across the seawater datasets, are examples of such a model (Giovannoni et al., 2005). Similarly, the higher energetic costs of production (Rocha & Danchin, 2002) and nitrogen requirement (McEwan et al., 1998) of guanine and cytosine, compared to adenine and thymine, may contribute to stronger selection for lower genomic GC content in seawater microbes.

Community profiles based on 876 seawater MAGs (Terzin et al., 2026) showed that sampling trip explained the largest proportion of community variation (58.6%), followed by reef site (23.8%) and GBR sector (9.7%). Applying the same sequential PERMANOVA to the 1,201 sediment MAGs, we instead identified reef site as the strongest driver (54.7%), followed by sampling trip (22.3%) and GBR sector (11.3%; Fig. S3a), consistent with the pattern observed in gene-centric mOTU profiles (Fig. 1g-h). Reef zoning status explained only a small proportion of the variation in both sediment (1.0%, *p* < 0.001) and seawater (1.9%, *p* < 0.001; Terzin et al., 2026) communities. Unlike seawater MAGs, which effectively distinguished no-take marine reserves and fished reefs (MINT sPLS-DA balanced accuracy 70.4%, *p* = 0.001; Terzin et al., 2026), sediment MAGs showed no predictive power (balanced accuracy 53.7%, *p* = 0.346; Fig. S3b-c). Gene-centric mOTU profiles yielded a similar result (balanced accuracy 51.2%, *p* = 0.409; Fig. S3d), suggesting that the absence of a zoning signal is unlikely to reflect incomplete MAG representation.

Reef zoning has been proposed to influence planktonic microbes through changes in fishing pressure and associated nutrient dynamics (Terzin et al., 2026). Efficient nutrient uptake and cycling within no-take reefs (high fish biomass, herbivory, and coral cover) may select for oligotrophic, streamlined bacterioplankton, whereas more nutrient-rich conditions within fished reefs (higher turf-to-coral ratios) may favour copiotrophic microbes with larger genomes (Terzin et al., 2026). We found no comparable effect of zoning on sediment microbial communities, which may be explained by the already nutrient-enriched nature of reef sediments (Entsch et al., 1983; Rasheed et al., 2002) and the rapid assimilation of any zoning-related nutrient enrichment by plankton before it reaches the benthic environment (Furnas et al., 2005; Sorokin, 1990). Rather, reef site explained the largest proportion of variation in sediment microbial communities, likely reflecting local differences in sediment properties (Glasl et al., 2019; Olmos Pin et al., 2026). This strong site effect may have masked the influence of other environmental factors, including reef zoning.

### Sediment microbes exhibit broad metabolic and respiratory versatility

To examine whether the observed differences in genomic architecture and diversity between habitats translate into functional divergence, we adopted a genome-centric approach based on genes encoded in the recovered MAGs, and a gene-centric approach based on predicted genes from all assembled contigs. In the genome-centric approach, we analysed 796 MAGs with ≥ 85% completeness (347 from the sediment, 449 from seawater). Comparative analysis revealed significant enrichment (Mann-Whitney U-test, adjusted *p* ≤ 0.05) of 1,224 KOs, 275 CAZymes, 3,835 Pfams, 870 peptidases, and 20 sulphatases in sediment MAGs relative to seawater MAGs (Tables S10 to S14). In comparison, seawater MAGs were enriched in a different set of 289 KOs, 103 CAZymes, 898 Pfams, 267 peptidases, and 10 sulphatases (Tables S10 to S14). The greater number of enriched features in sediment MAGs is partly expected given their larger genomes (approximately 50% larger than seawater MAGs), encoding more genes overall. Nonetheless, these results indicate greater functional diversity and complexity among reef sediment microbes.

Similarly, we examined the distribution of 44 signature metabolic marker genes and 30 hydrogenase subgroups (Materials and Methods; Tables S15 and S16), using genome-centric (Fig. 3a-b) and gene-centric (Fig. 3c) approaches. Across both sediment and seawater microbes, genes associated with aerobic respiration were broadly detected, including ATP synthase (*atpA*), NADH:quinone oxidoreductase (*nuoF*), succinate dehydrogenase (*sdhA*), and cytochrome *c* oxidase subunits (*coxA* and *cyoA*). However, sediment microbes were significantly enriched in high-affinity terminal oxidases, including cytochrome *bd* ubiquinol oxidase (*cydA*) and *cbb_3_*-type cytochrome c oxidase (*ccoN*), alongside genes for respiration using alternative electron acceptors and donors (e.g. *frdA* for fumarate reduction, *rdhA* for reductive dehalogenation, and *fdhA* for formate oxidation). Furthermore, we infer greater capacity for dissimilatory nitrogen and sulphur respiration among the sediment microbes than their seawater counterparts (Fig. 3a-c). For nitrogen, sediment microbes were enriched in the full denitrification pathway of nitrate reduction (*narG* and *napA*; NO_3_^−^ to NO_2_^−^), nitrite reduction (*nirKS*; NO_2_^−^ to NO), nitric oxide reduction (*norB*; NO to N_2_O), and nitrous oxide reduction (*nosZ*; N_2_O to N_2_), alongside the marker gene (*nrfA*) for dissimilatory nitrate reduction to ammonium (DNRA; NO_2_^−^ to NH_4_^+^). For sulphur, sediment microbes were enriched in genes encoding dissimilatory sulphate reduction (*dsrA*), sulphide oxidation (*sqr* and flavocytochrome *c* sulphide dehydrogenase, FCC), sulphur disproportionation (*sor*), and thiosulphate oxidation (*soxB*). These results suggest that sediment microbes are equipped with a continuum of aerobic, microaerobic, and anaerobic respiratory capacity, likely reflecting the irregular fluctuations in redox conditions at surface layers of permeable marine sediments (Cook et al., 2007; Huettel et al., 2014). Such flexibility in energy-yielding mechanisms was largely underrepresented in aerobically respiring seawater microbes.

**Fig. 3.**
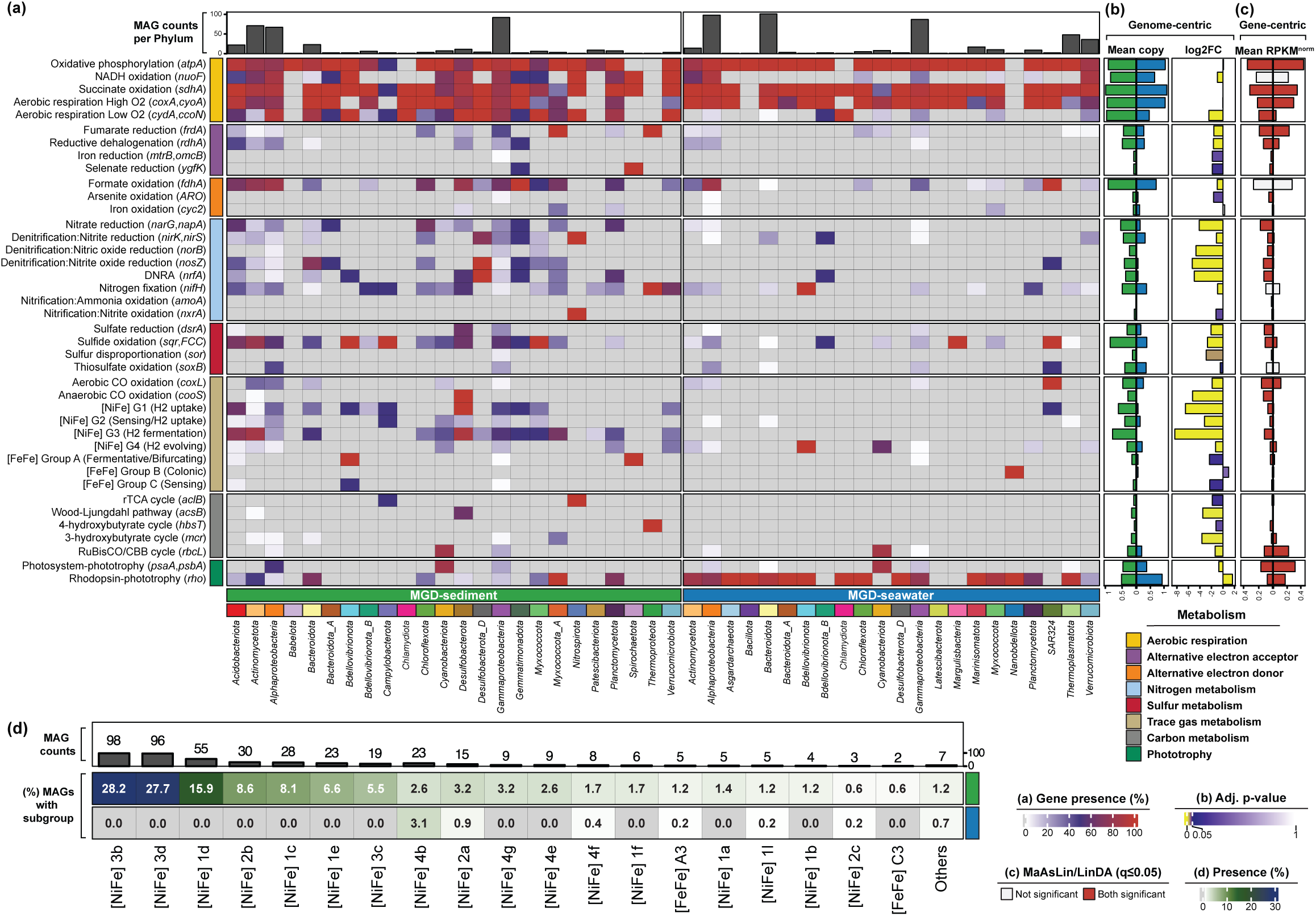
**Metabolic capacity of sediment and seawater microbes of the Great Barrier Reef. (**a) Heatmap showing the distribution of key metabolic marker genes across metagenome-assembled genomes (MAGs) with completeness ≥ 85% from sediment (347 MAGs; left) and seawater (449 MAGs; right), displayed as the proportion of MAGs encoding each gene within each phylum (*Pseudomonadota* split into *Alphaproteobacteria* and *Gammaproteobacteria*). The top bar plot indicates the number of MAGs per phylum (or class for *Pseudomonadota*). (b) Side panels showing results from the genome-centric analysis, from left to right: mean copy number of each gene in sediment (left) and seawater (right) MAGs; log_2_ fold-changes in mean copy number of each gene, where negative values indicate enrichment in sediment and positive values indicate enrichment in seawater. Colours reflect adjusted *p*-values from the Benjamini-Hochberg procedure. (c) Results from the gene-centric analysis, displaying mean of normalised reads per kilobase per million (RPKM) values for each gene in sediment (left) and seawater (right) communities. Bars are coloured red if statistical significance (adjusted *p* ≤ 0.05) was supported by both MaAsLin2 and LinDA. (d) Heatmap showing the fraction of sediment and seawater MAGs encoding each hydrogenase subgroup as per the HydDB classification. The top bar plot shows the total number of MAGs encoding each subgroup.

### Sediment microbes exhibit potential for nitrification and nitrogen fixation

Nitrification is a critical process that mitigates nitrogen loss in reef ecosystems, commonly initiated with ammonia monooxygenase (encoded by *amoA*). Our gene-centric analysis revealed higher abundance of *amoA* genes in sediment communities (Fig. 3c), with all *amoA* sequences classified as archaeal type based on their phylogenetic placement (Materials and Methods; Table S17): this was consistent with the detection of *amoA* only in *Nitrosopumilaceae* (4 MAGs; Table S18) across all sediment and seawater MAGs. Nitrite oxidation is the subsequent step involving nitrite oxidoreductase (*nxrA*), which was enriched in the sediment microbes (Fig. 3b-c) and detected in a sediment *Nitrospiria* MAG. Collectively, these results suggest that ammonia-oxidising archaea, particularly *Nitrosopumilaceae*, are key contributors to ammonia oxidation and nitrogen cycling in reef sediments. This pattern is consistent with earlier studies implicating *Nitrosopumilus* in ammonia oxidation in permeable marine sediments (Gaidos et al., 2011), due to their higher affinity for NH_4_^+^ under low concentrations (Martens-Habbena et al., 2009; Rusch & Gaidos, 2013; Wang et al., 2017).

Beyond nitrification, a greater diversity of microbes with the potential for nitrogen fixation was found in sediments, compared to their seawater counterparts (Fig. 3a-c), with the catalytic component of nitrogenase (*nifH*) detected in 75 sediment MAGs representing 15 phyla (e.g. *Pseudomonadota*: 28 MAGs, *Actinomycetota*: 15, Acidobacteriota: 9), compared to 54 seawater MAGs representing 8 phyla. These findings suggest substantial capacity for biological nitrogen fixation in reef sediments. Nitrogen fixation that converts gaseous N_2_ to bioavailable NH_4_^+^ is a key process alleviating nitrogen limitation in coral reef ecosystems (Cardini et al., 2016), where bioavailable nitrogen is often scarce (Furnas et al., 2011). Although N_2_ fixation rates in surface sediments are considered lower than those of other benthic substrates (e.g. microbial mats and seagrass beds), sediment microbes may still contribute substantially to reef nitrogen budgets because of their extensive spatial coverage (O’Neil & Capone, 2008).

### Trace gas metabolism as a redox-balancing mechanism in sediment microbes

The potential for trace gas metabolism emerged as a key distinguishing feature of sediment microbes. Sediment MAGs were substantially enriched with genes encoding both aerobic and anaerobic oxidation of carbon monoxide (CO; *coxL* and *cooS*) and nearly all detected hydrogenase subgroups: hydrogenases were detected in 66% (230 of 347) of sediment MAGs and in only 6% (26 of 449) of seawater MAGs. Nickel-iron [NiFe]-hydrogenases were prevalent in both habitats (227 sediment MAGs and 24 seawater MAGs), whereas [FeFe]-hydrogenases were rare (6 sediment MAGs and 2 seawater MAGs), and [Fe]-hydrogenases were absent (Fig. 3d). The two most abundant hydrogenase subgroups in sediment MAGs were the cofactor-coupled bidirectional [NiFe] 3b (98 MAGs) and 3d (96 MAGs; Fig. 3d), which can couple NAD(P)H oxidation to H_2_ production (i.e. hydrogenogenic) to maintain cellular redox balance; these subgroups were absent from seawater MAGs. Of the 172 sediment MAGs encoding [NiFe] 3b and/or 3d (22 encode both), 91 also encode terminal oxidases for aerobic respiration (e.g. *coxA*, *ccoN*, and *cydA*; Table S16). Sediment MAGs also encode H_2_-consuming [NiFe] hydrogenases that mediate aerobic ([NiFe] 1d; 55 MAGs) and anaerobic ([NiFe] 1c; 28) hydrogenotrophic respiration, as well as subgroups involved in H_2_-sensing ([NiFe] 2b), electron bifurcation from H_2_ to heterodisulphide ([NiFe] 3c), and hydrogenotrophic respiration using sulphur ([NiFe] 1e) or sulphate and other electron acceptors ([NiFe] 1a and 1b). The potential for both hydrogenogenic fermentation and hydrogenotrophic respiration was observed in sediment MAGs representing several highly prevalent families (> 80% of all samples), including *Rhodobacteraceae*, *Woeseiaceae*, *SZUA-36*, *SZUA-38*, *UBA5794*, and *Ilumatobacteraceae* (Tables S5 and S16). These results suggest that hydrogen metabolism is an ecologically significant feature of reef sediment microbes, acting as a flexible redox valve that buffers transient imbalances in intracellular reducing equivalents (Berney et al., 2014). The prevalence of sediment MAGs encoding both fermentative and respiratory hydrogenases indicates that some taxa can switch between H_2_ production during anoxia and H_2_ consumption upon reoxygenation, a strategy previously reported for facultative anaerobes inhabiting coastal surface sediments (Chen et al., 2021; Kessler et al., 2019). Such flexibility may help sediment microbes maintain activity as coral reefs experience more frequent hypoxic events associated with ocean warming, deoxygenation, and reef degradation (Altieri et al., 2017; Hughes et al., 2020; Pezner et al., 2023). Recurrent hypoxia may also alter the balance between production and consumption of H_2_ in reef sediments. Given the significance of H_2_ as an indirect greenhouse gas (Derwent et al., 2006), determining whether reef sediments act as net sources or sinks of H_2_ warrants investigation using flux measurements. Transcriptomic and enzymatic data will further be needed to determine whether hydrogenogenic and hydrogenotrophic capacities are actively expressed or functionally uncoupled in reef sediments.

The [NiFe]-hydrogenases commonly associated with aerobic H_2_ uptake (subgroups 1d, 1l, and 2a) were absent or rare among reef seawater MAGs (0%, 0.2%, and 0.9%; Fig. 3d), indicating a limited potential for aerobic H_2_ oxidation. In support of this, some of the dominant reef water taxa (e.g. *Prochlorococcus* and *Synechococcus*) are reported to lack hydrogenase genes, potentially driven by genome streamlining (Barz et al., 2010). [NiFe] 4b hydrogenases, typically associated with formate or CO dehydrogenase complexes, were more prevalent among seawater MAGs (14) than in sediment MAGs (9). Together with the moderate abundance of *coxL* genes in seawater MAGs and communities (Fig. 3a-c), these results suggest CO instead of H_2_ as a more broadly utilised trace gas supporting microbial metabolism in reef surface waters. The abundance of phototrophic taxa in well-lit surface waters likely favours harvest of solar energy at lower metabolic cost than H_2_ metabolism that requires specialised metalloenzymes (Greening & Grinter, 2022; Lappan et al., 2023). Consistent with this interpretation, the capacity for bacterial H_2_ oxidation is known to increase with depth and in marine regions or seasons characterised by reduced primary production (Lappan et al., 2023).

### Sediment and seawater microbes exhibit diverging strategies in carbon cycling, polysaccharide utilisation, and phototrophy

Transitions between fermentative and respiratory modes in sediment microbes are likely sustained by access to stored and recycled carbon pools. Consistent with this concept, sediment microbes exhibited enrichment in genes encoding the reductive tricarboxylic acid (rTCA) cycle (*aclB*), reductive acetyl-CoA (Wood-Ljungdahl; WL) pathway (*cooS*, *acsB*), 4-hydroxybutyrate cycle (*hbsT*), and 3-hydroxypropionate cycle (*mcr*). Although the gene coding for Calvin-Benson-Bassham (CBB) cycle (*rbcL*; RuBisCO) was more frequently detected among sediment MAGs, this gene was more abundant in seawater communities (gene-centric analysis; Fig. 3c). In parallel, genes for rhodopsin (*rho*)-and photosystem-based light harvesting (*psaA* and *psbA*) were enriched in seawaters (Fig. 3a-c), reflecting the prevalence of photoautotrophic carbon fixation in well-lit and oxygen-rich surface waters (Flombaum et al., 2013). These results suggest that, compared to seawater microbes, sediment microbes may rely less on continuous photoautotrophic carbon fixation, and more on chemolithotrophic and carbon-recycling strategies to support their activity in a light-limited environment.

To identify putative sources and processing strategies of organic carbon in sediment and seawater microbes, comparative enrichment analyses of CAZymes, peptidases, and sulphatases were conducted using a genome-centric approach on the 796 MAGs with ≥ 85% completeness (Materials and Methods; Table S19). Sediment MAGs were enriched with 275 CAZymes, including four auxiliary activities (AAs), 19 carbohydrate-binding modules (CBMs), 15 carbohydrate esterases (CEs), 113 glycoside hydrolases (GHs), 122 glycosyltransferases (GTs), and two polysaccharide lyases (PLs). In contrast, seawater MAGs exhibited narrower functional enrichment: 103 CAZymes comprising 10 CBMs, eight CEs, 41 GHs, and 44 GTs. In total, the identified CAZymes formed 20,259 CAZyme Gene Clusters (CGCs; 10,040 from 347 sediment MAGs, 10,219 from 449 seawater MAGs), of which only a small subset (1,678 CGCs; 750 from sediment MAGs, 928 from seawater MAGs) have substrate predictions.

Organic particles settle from the water column to the seafloor, and the preferential use of simple and soluble carbohydrates by seawater microbes leaves recalcitrant compounds only partially degraded, likely leading to their accumulation in benthic environments (Viver et al., 2025). Our results support this notion; sediment MAGs were enriched in GH36 α-galactosidase, known for its activity on α-galactan/raffinose (Liebl et al., 1998; Reddy et al., 2016), and β-mannanase (GH113) involved in β-mannan degradation (Fig. 4a; Couturier et al., 2022). Together with pathways that process other low-solubility substrates (i.e. alginate and pectin), CGCs targeting complex carbohydrates were more abundant among sediment MAGs (Fig. 4b). Sediment MAGs were also enriched in CAZymes associated with the catabolism and modification of peptidoglycan, a substrate often considered recalcitrant (Jørgensen et al., 2003). Significant enrichment of GT51 and GT28 suggests a greater potential for peptidoglycan biosynthesis, whereas enrichment of GH179, GH23, GH73, GH102, GH103, CE4, CE14, and peptidoglycan recognition module CBM50 indicates a capacity for peptidoglycan turnover (Fig. 4a). Complementing this pattern, diverse MEROPS peptidase gene families (870) were enriched in the sediment MAGs relative to seawater MAGs (267), highlighting the proteolytic potential of sediment microbes to degrade complex proteinaceous biomass (Du et al., 2016).

**Fig. 4.**
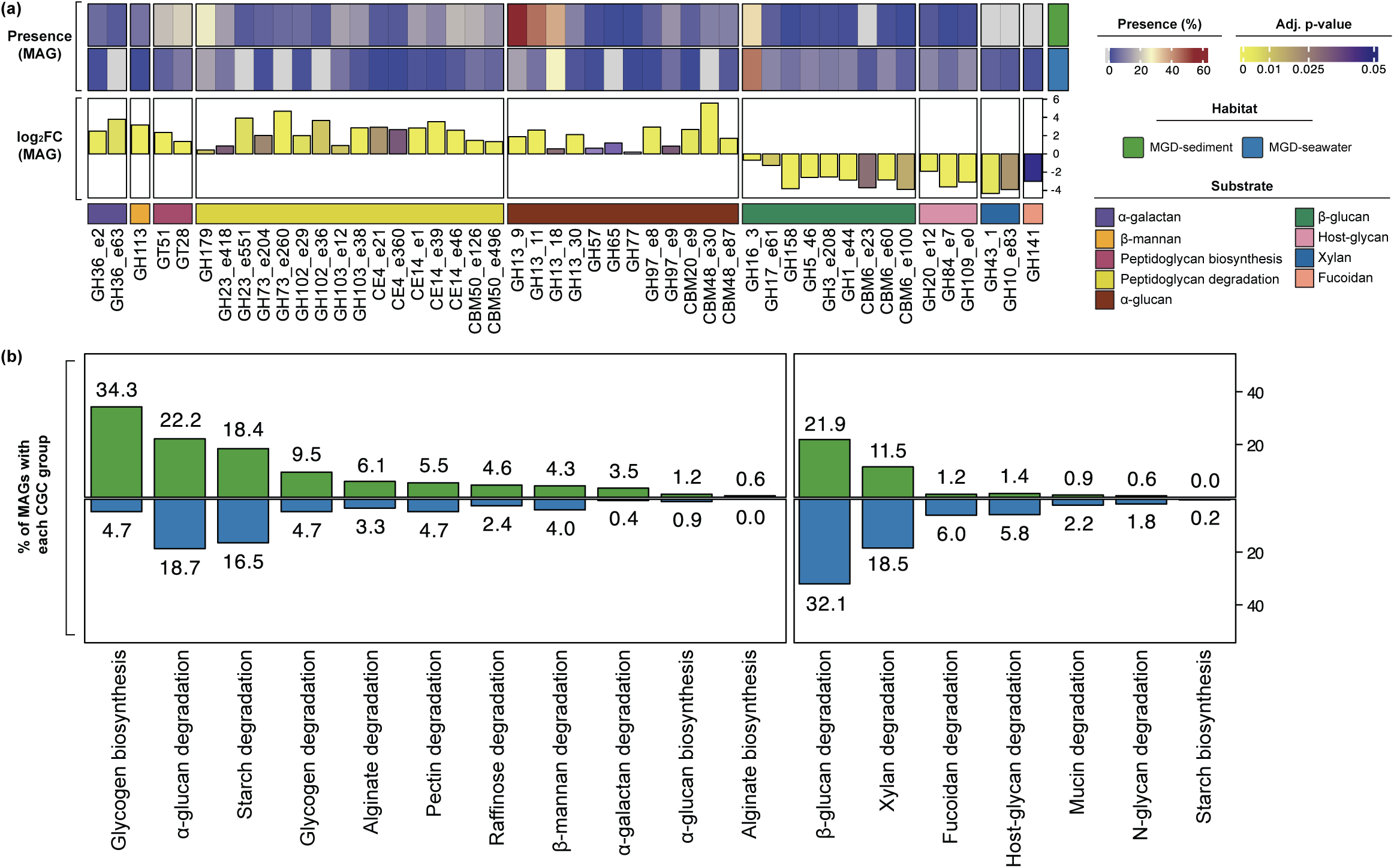
**Potential for polysaccharide biosynthesis and degradation in sediment and seawater microbes of the Great Barrier Reef. (**a) Heatmap showing the distribution of key CAZymes across metagenome-assembled genomes (MAGs) with completeness ≥ 85% from sediment (347 MAGs) and seawater (449 MAGs), displayed as the proportion of MAGs encoding each CAZyme within each habitat. The bar plot below displays log_2_ fold-changes in mean copy number of each CAZyme, where positive values indicate enrichment in sediment and negative values indicate enrichment in seawater. Colours reflect adjusted *p*-values from the Benjamini-Hochberg procedure. (b) Bar plot showing the fraction of sediment (green; top) and seawater (cyan; bottom) MAGs encoding each CAZyme gene cluster group (defined by substrate target and mode of action as per dbCAN-PUL).

Despite their high solubility and structural simplicity, CAZymes involved in degrading α-glucans, the common storage compounds in marine organisms (e.g. algae, animals, and bacteria), were significantly enriched in sediment MAGs. These include GH13, GH57 (associated with type III α-glucan PULs; Wang et al., 2024), GH65 (type II; Kruger et al., 2019), GH77 (type III), GH97 (type II), and starch-binding modules of CBM20 and CBM48 (Fig. 4a). Similarly, sediment MAGs exhibited greater prevalence of CGCs for degrading starch, glycogen, and α-glucans. Whereas biosynthetic potential for starch and α-glucans was largely similar between sediment and seawater MAGs, the most substantial difference was observed for CGCs driving glycogen biosynthesis, detected in 34.3% of sediment MAGs compared to 4.7% of seawater MAGs (Fig. 4b). These CGCs were broadly distributed across bacterial phyla (Tables S16 and S20), including *Gammaproteobacteria* (56 MAGs), *Actinomycetota* (32), *Alphaproteobacteria* (23), and *Acidobacteriota* (7). Bacterial glycogen synthesis is a survival strategy that preserves cell integrity and enables adaptation to starvation and stress (Sekar et al., 2020; Wang et al., 2020). Glycogen likely serves as a key energy reserve for microbes in reef sediments; it can be synthesised during high carbon influx and endogenously mobilised during transient anoxia or limiting terminal electron acceptors, to sustain fermentative metabolism and nitrate/sulphate reduction, thereby maintaining ATP production and redox balance (McIlroy et al., 2014). Glycogen biosynthesis CGCs were frequently detected in MAGs belonging to *Woeseiaceae* (10 of 12 high-completeness MAGs) and *SZUA-36* (15 of 18 high-completeness MAGs), two highly abundant and prevalent gammaproteobacterial families across our sediment samples (Tables S5 and S16), underscoring their capacity to cope with environmental heterogeneity in reef sediments.

These enrichment patterns also suggest the capacity of sediment microbes to exploit organic matter fuelled by dead microbial cells (i.e. necromass), such as peptidoglycan from cell walls and intracellular α-glucans. Degradation rates of extracellular DNA, a proxy commonly used to estimate necromass turnover in terrestrial soils (Ni et al., 2020), are known to be higher in marine benthos compared to the water column (Dell’Anno & Corinaldesi, 2004), highlighting the importance of necromass turnover in sediment environments (Orsi et al., 2018; Pelikan et al., 2021). We observed co-enrichment of biosynthetic and degradation pathways for necromass-associated biopolymers among the sediment MAGs; this suggests an intra-population recycling loop, similar to that proposed for bacterioplankton and phytoplankton blooms in the water column (Beidler et al., 2024). Further investigation of temporal and seasonal variations will clarify the source and relative importance of these substrates in sedimentary environments.

Seawater MAGs were enriched with CAZymes (e.g. GH16_3, GH17, GH158, GH5_46) associated with PULs targeting β-glucan and laminarin (Kruger et al., 2019; Sidhu et al., 2023; Wang et al., 2024). Laminarin, a water-soluble β-1,3-glucan with β-1,6 branches, is the main storage carbohydrate of phytoplankton, brown macroalgae, and other primary producers, and is one of the most important sources of organic carbon in the ocean (Becker et al., 2020). Enrichment was also observed for the CBM6 domain, previously reported to bind β-1,3/1,4-glucans (Michel et al., 2009). Consistent with this, 210 β-glucan-degrading CGCs were detected in 32.1% of seawater MAGs, compared to 98 in 21.9% of sediment MAGs (Fig. 4b). Many of these β-glucan-degrading CGCs (243 of 308) resembled PULs that were experimentally characterised to degrade laminarin (Table S20). Of these 243 putative laminarin-degrading CGCs (Table S20), 120 were recovered from *Bacteroidota*, predominantly the order *Flavobacteriales* (107 CGCs) and family *Flavobacteriaceae* (60 CGCs). These findings are consistent with the recognised role of *Flavobacteriaceae* in the transformation of marine organic carbon, including laminarin, into CO_2_ or biomass, especially during phytoplankton blooms (Buchan et al., 2014). Additionally, 169 of the 243 laminarin-degrading CGCs were recovered from seawater MAGs (Table S20), suggesting enhanced capacity for laminarin degradation that supports fast microbial growth and efficient carbon turnover (Kappelmann et al., 2019). This pattern likely reflects the ubiquity, solubility, and structural simplicity of laminarin (Becker et al., 2020). Since laminarin is produced in such large quantities by primary producers in marine environments, a substantial fraction is expected to reach marine sediments. This may explain the moderate abundance of GH16_3 (Fig. 4a) and β-glucan-targeting CGCs in sediment MAGs (Fig. 4b), suggesting that laminarin and other β-glucans are also important carbon sources for sediment microbes (Lindsay et al., 2025).

### Sediment microbes exhibit rich and distinctive secondary metabolite biosynthesis potential

We compared 3,504 BGCs identified from the MGD-sediment dataset with those from MGD-seawater (1,472), open ocean (OMD; 16,335), and reef invertebrates (RMD; 12,259). After clustering into non-redundant gene cluster families (GCFs; Tables S21 and S22), MGD-sediment showed the highest biosynthetic richness (2.6 GCFs per species), compared to those from MGD-seawater (1.3), OMD (1.4), and RMD (2.3; Fig. 5a). Compared to others, GCFs of MGD-sediment revealed a higher proportion of non-ribosomal peptide synthetase (NRPS) pathways and a lower fraction of terpene biosynthesis pathways (Fig. 5b). Many NRPS-derived peptides function as antibiotics, toxins, or siderophores, suppressing competitors or facilitating sequestration of limiting nutrients (Geller-McGrath et al., 2023; Miethke & Marahiel, 2007). NRPS pathways may be selected in crowded sedimentary environments, where direct antagonistic metabolites or efficient nutrient acquisition confer competitive advantages. This observation aligns with a previous study reporting the highest NRPS richness in microbes of marine sediments relative to those of terrestrial soil and seawater and suggesting that heterotrophic taxa in marine sediments employ NRPS pathways to compete for organic carbon resources (Geers et al., 2023). In contrast, terpenes, such as carotenoids, can act as antioxidants, photoprotective pigments, or membrane-stabilising compounds under light-induced stress (Steven et al., 2022). These functions are more advantageous in light-exposed pelagic environments and may explain the higher prevalence of terpene biosynthesis pathways among reef seawater and open ocean microbes.

**Fig. 5.**
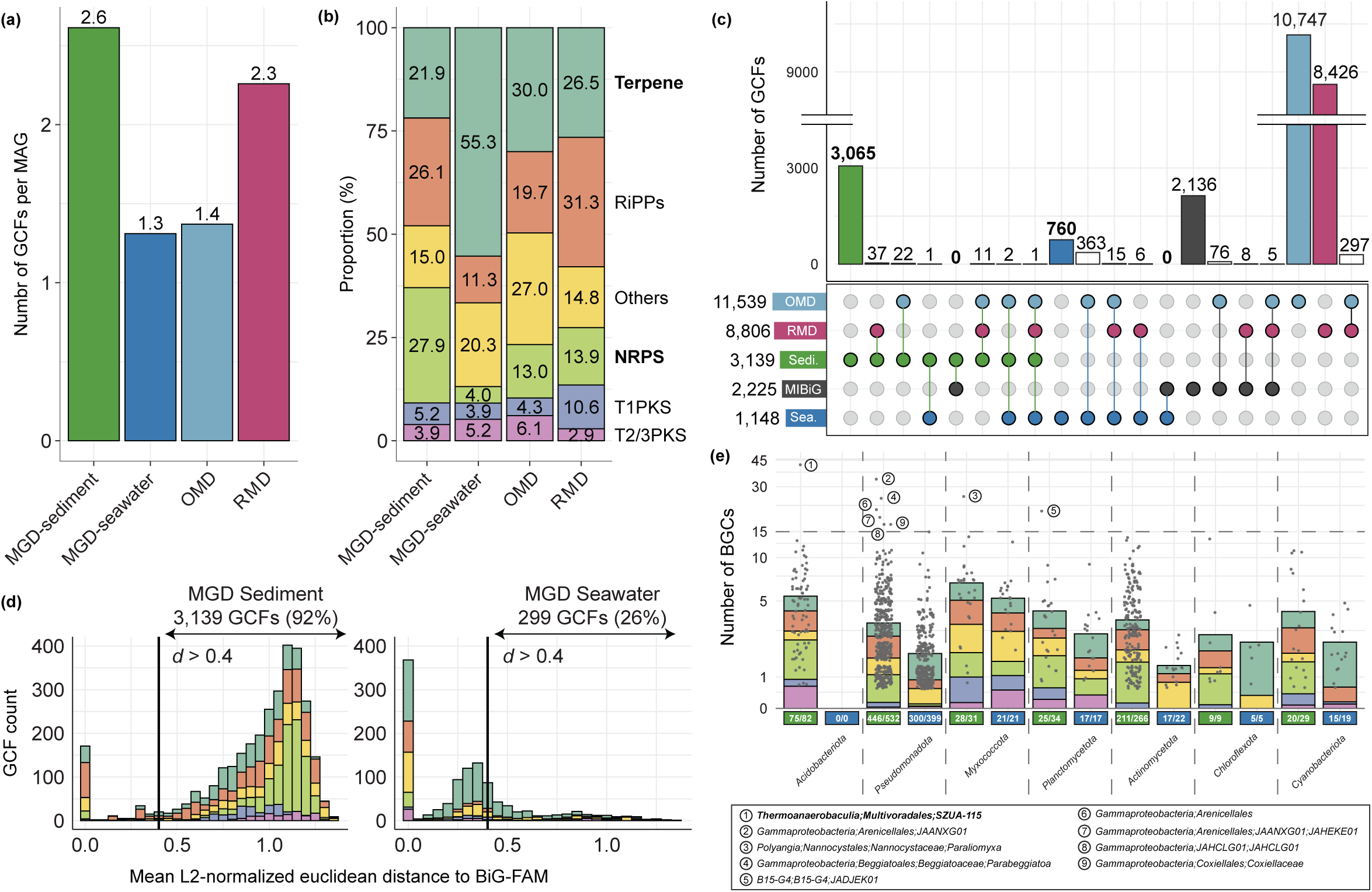
**Biosynthetic potential of sediment and seawater microbes of the Great Barrier Reef. (**a) Bar plot showing the number of gene cluster families (GCFs) per dereplicated metagenome-assembled genome (MAG) from reef sediment (MGD-sediment), reef seawater (MGD-seawater), the open ocean (Ocean Microbiomics Database; OMD), and reef-associated invertebrates (Reef Microbiomics Database; RMD). (b) Composition of natural product classes encoded by GCFs in each dataset. (c) UpSet plot showing the specificity and novelty of MGD-sediment GCFs, compared with biochemically characterised pathways in the MIBiG database and with predicted pathways from the other datasets. (d) Mean L2-normalised Euclidean distance between GCFs from MGD-sediment (left) and MGD-seawater (right) and biosynthetic gene clusters (BGCs) predicted from genomes deposited in RefSeq (BiG-FAM). The vertical line indicates the novelty threshold (*d* = 0.4). (e) Scatter plot showing the number of BGCs detected in sediment and seawater MAGs from seven phyla, each containing at least one sediment MAG encoding more than ten BGCs. Nine BGC-rich species, defined as MAGs encoding more than 15 BGCs, are labelled numerically. The overlaid stacked bar plot shows the mean number of BGCs per natural product class in each phylum. The fraction of MAGs with BGCs in each phylum is shown at the bottom of each bar.

GCFs of MGD-sediment were highly distinctive, with only 74 (2.4% of 3,139; Fig. 5c) shared with other marine microbes; this compares to 388 (33.8% of 1,148) for MGD-seawater. While a subset of GCFs from OMD (76) and RMD (8) overlapped with pathways in the MIBiG database (Fig. 5c), no GCFs from either MGD dataset intersected with this catalogue. When compared to pathways predicted from ∼190,000 genomes deposited in RefSeq (BiG-FAM; Fig. 5d), 92% of the GCFs predicted from MGD-sediment were classified as novel (*d* > 0.4), compared with 26% for MGD-seawater, underscoring a higher degree of novelty and/or habitat specificity of biosynthetic repertoires in reef sediments.

To facilitate future experimental characterisation of these putatively novel pathways among biosynthetically rich taxa, we focused on MAGs encoding > 15 BGCs. Whereas no seawater MAGs met this criterion, nine candidate BGC-rich species were identified from sediment MAGs (Fig. 5e), all of which are new candidate species representing the phyla *Myxococcota*, *Acidobacteriota*, *Planctomycetota*, and *Pseudomonadota*. The most BGC-rich lineage belongs to the order *Multivoradales* (formerly *Ca. UBA5704*; class *Thermoanaerobaculia*), supporting the notion that marine *Acidobacteriota* and the other three phyla are key reservoirs of biotechnologically relevant biosynthetic potential (Gregory et al., 2019; Leopold-Messer et al., 2023; Paoli et al., 2022; Timmermans et al., 2017; Wiederkehr et al., 2026; Wiegand et al., 2020).

Together, these results demonstrate that reef sediment microbes harbour exceptionally rich, highly novel, and functionally specialised biosynthetic repertoires. This pattern likely reflects strong selection for secondary metabolites to enhance fitness in a dense, resource-variable environment, through direct chemical interference and efficient nutrient acquisition, driving niche differentiation (Junkins et al., 2022; Penn et al., 2009). Earlier genome-mining efforts in the open ocean (Paoli et al., 2022) and in coral and sponge microbiomes (Wiederkehr et al., 2026) revealed extensive, largely uncharacterised biosynthetic potential and led to the discovery of new enzymes and natural products. Complementing these pelagic and host-associated surveys, our results highlight reef sediments as a major, underexplored source of new biosynthetic enzymes and natural products. Experimental characterisation of the predicted BGCs is essential to resolve the ecological roles of these metabolites in shaping microbial fitness and community dynamics.

## Conclusions

In this study, we characterised microbial communities inhabiting surface sediments of the GBR using gene-and genome-centric metagenomics, and in the process have established the most extensive genomic reference of reef sediment microbes to date. Based on extensive taxonomic, metabolic, and biosynthetic novelty revealed from these data, our results highlight reef sediments as a major, underexplored reservoir of microbial diversity with significant potential for nutrient cycling and natural product discovery, likely driven by frequent yet irregular spatiotemporal variation in redox conditions and nutrient availability. By contrast, seawater communities were dominated by taxa adapted to oligotrophic and oxygen-rich conditions. This niche differentiation reflects opposing selection regimes of metabolic flexibility versus growth efficiency, highlighting how environmental heterogeneity and resource availability shape microbial evolution. As these contrasting communities co-occur at the same sites and times, they may play complementary roles in reef nutrient cycling through benthic-pelagic coupling, underscoring the need to incorporate benthic microbial processes into models of coral reef ecology and function. Future efforts integrating metagenomics with metatranscriptomics, metabolomics, and process-rate measurements of microbial activity will be critical to validate genomic predictions and assess their ecological significance under changing reef conditions.

## Supporting information

Supplementary Figures S1 through S3

Supplementary Tables S1 through S22

## Acknowledgements

The sediment samples in this study were collected across 48 sites from the sea country of multiple Indigenous groups who are Traditional Owners (TOs) of that land and sea. We acknowledge the Gudang Yadhaigana TOs, custodians of the McSweeney, Monsoon, 11-049, and 11-162 reefs, which lie within their sea country estate. We pay our respects to the Kuuku Ya’u TOs of the Mantis and Lagoon reefs, and the Lama Lama TOs of the 13-124, Davie, and Corbett reefs in the western half of their territory. We also recognise the Cape Melville, Howicks, and Flinders Island TOs of the eastern half of Corbett and Sand bank #1 reefs, as well as the Eastern Kuku Yalanji TOs of St Crispin and Agincourt #1. We extend our respect to the Yirrgandji TOs of Hastings reef and the Gunggandji as TOs of Arlington, Thetford, and Moore reefs. We acknowledge the Gunggandji-Mandingalbay Yidinji TOs of McCulloch, Hedley, Peart, and Feather reefs, and the Mandubarra TOs of Farquharson reef. We honour the Girringun Aboriginal Corporation TUMRA for their connection to Taylor reef, and the Manbarra TOs of Rib, Kelso, Little Kelso, and John Brewer reefs. We also recognise the Wulgurukaba TOs of Myrmidon, Grub, and Helix reefs, the Bindal TOs of Knife, Fork, Centipede, Chicken, and Lynchs reefs, and the continuing connection of the Manbarra TOs to Roxburgh, and Fore and Aft reefs. We acknowledge the PCCC TUMRA for their stewardship of North, Broomfield, Erskine, Masthead, Hoskyn, Fairfax, and Boult reefs. We pay our respects to their Elders, past, present, and emerging, and acknowledge their enduring connection to land and sea.

Part of this research took place at the Australian Institute of Marine Science (AIMS) headquarters at Cape Ferguson, and we wish to acknowledge the Wulgurukaba and Bindal peoples as the TOs of that land. This research was also undertaken at the James Cook University Townsville Bebegu Yumba campus. This research was also conducted at The University of Queensland St Lucia campus (Meanjin, Brisbane), and we acknowledge the Turrbal and Jagera peoples as the TOs of the lands. Authors acknowledge that the Australian Aboriginal and Torres Strait Islander peoples are the original inhabitants and traditional custodians of this continent and have unique cultural and spiritual relationships to the land and waters.

We acknowledge the AIMS-LTMP team and others involved in field collection and preparation of samples including Michael Emslie, Emmanuelle Botté, Johnston Davidson, Véronique Mocellin, Josephine Nielsen, and Christian O’Dea. We thank the crew of the RV Solander and RV Cape Ferguson for their excellent logistical support in the field.

## Data availability

Raw sequencing data will be available upon publication through the AIMS data repository (https://doi.org/10.25845/kkma-n127) and NCBI BioProject accession PRJEB112104. Sediment MAGs generated from this study are available at https://doi.org/10.5281/zenodo.24187765. Detailed methods, including additional data and code to replicate the analysis, are available at https://github.com/hisatakeishida/IMOS-GBR-Sediment-Microbe.

## Ethics approval and consent to participate

Samples were collected under the permit G12/35236-1 issued by the Great Barrier Reef Marine Park Authority.

