## Supplementary Figures S1 through S3 for "Comparative metagenomics reveals diversity and metabolic heterogeneity of sediment microbes from the Great Barrier Reef"

This supplementary material contains:

**Supplementary Figures S1 through S3**

**Supplementary Tables S1 through S22 (in a separate file)**

### Supplementary Figures

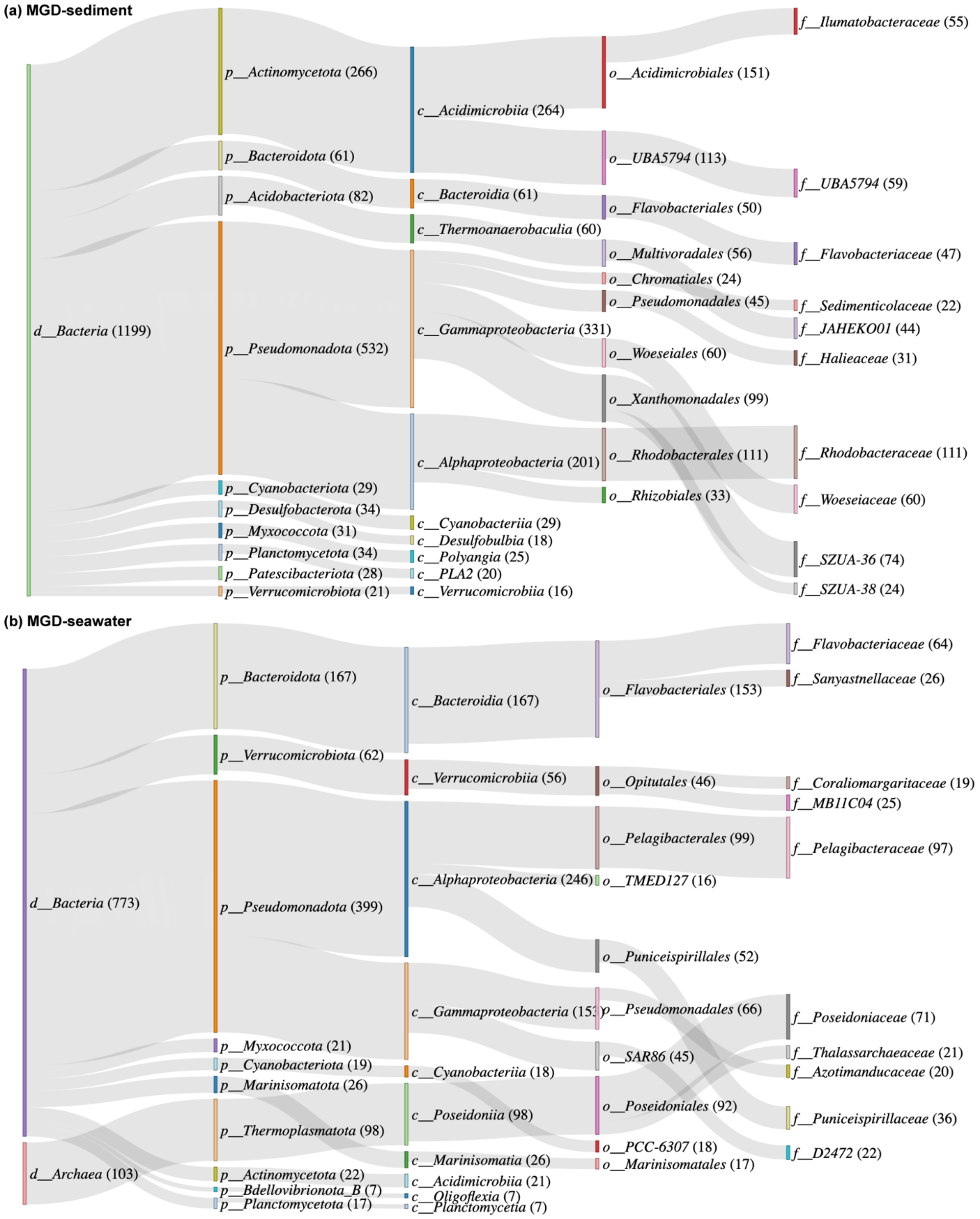


**Fig. S1 | Taxonomic assignment of metagenome-assembled genomes (MAGs) reconstructed from sediment and seawater samples of the Great Barrier Reef.**

Sankey diagram showing the taxonomic assignment down to the family level of (a) 1,201 sediment MAGs and (b) 876 seawater MAGs. Only the 10 most represented taxa at each rank are shown. The numbers of MAGs recovered for each taxon are indicated.

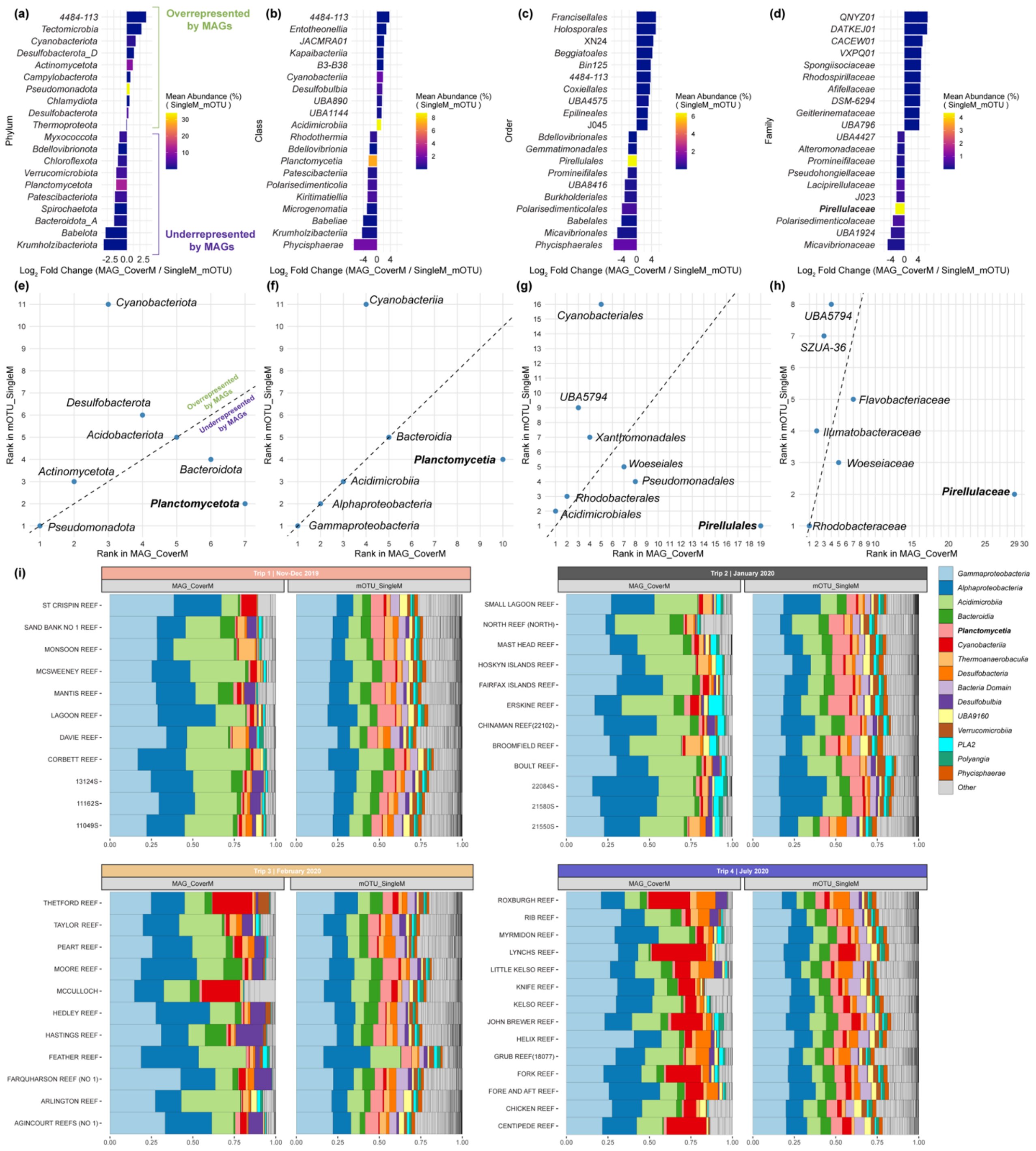


**Fig. S2 | Differences in sediment community composition captured by gene- and genome-centric approaches.**

(a-d) $\log_2$ fold-changes comparing relative abundance of microbial taxa in sediment communities of the Great Barrier Reef, estimated by gene- (SingleM) and genome-centric (CoverM) approaches at the phylum (a), class (b), order (c), and family (d) levels. Negative values indicate underrepresentation of taxa by CoverM-based profiles relative to SingleM-based profiles. (e-h) Comparison of the most abundant (top 5) prokaryotic taxa detected by SingleM (y-axis) and CoverM (x-axis) at the phylum (e), class (f), order (g), and family (h) levels. (i) Community composition at the class level (top 15), split by trip name (Trip1, Trip2, Trip3, Trip4) and method (CoverM on left and SingleM on right).

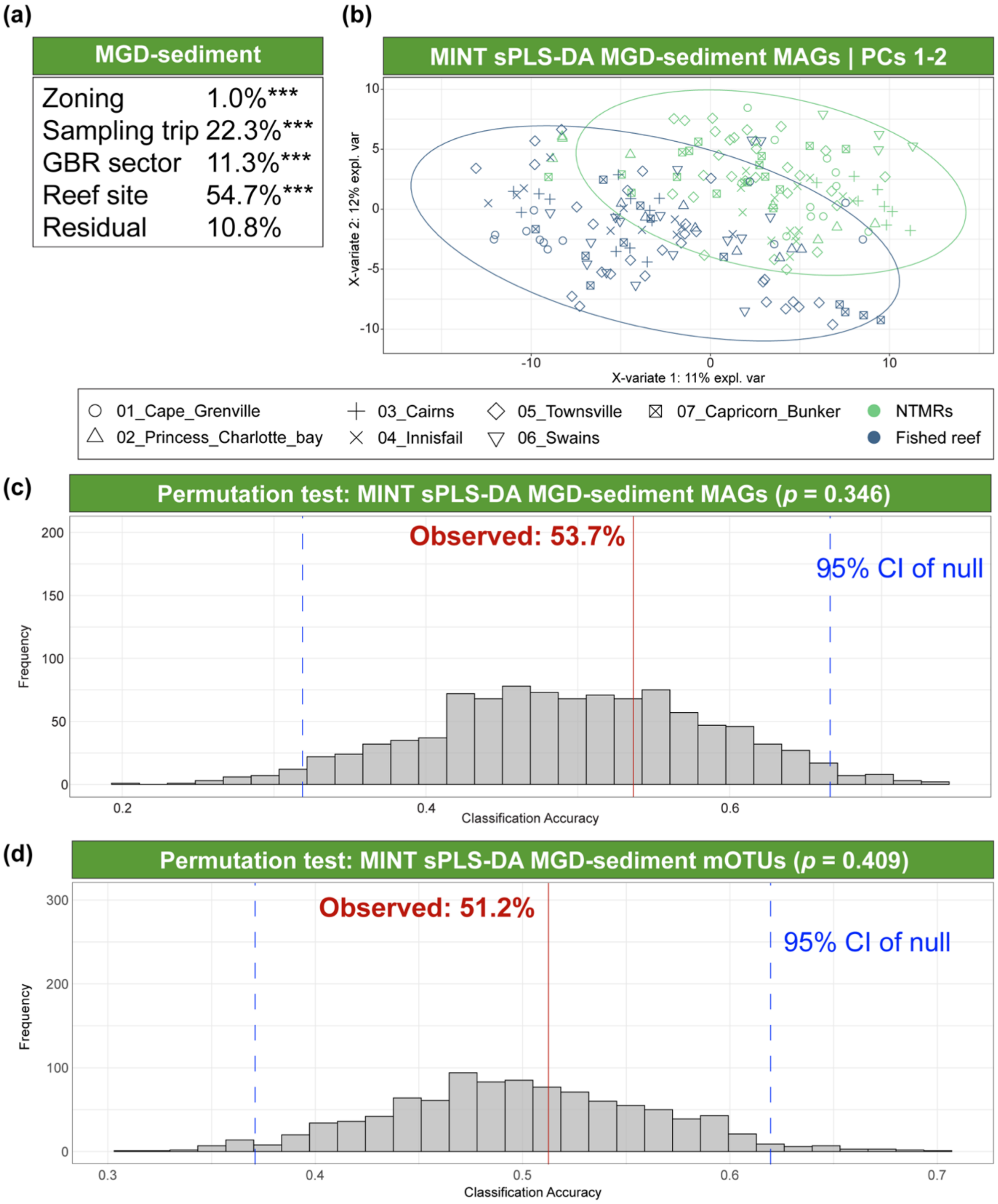


**Fig. S3 | Predicting reef zoning status using sediment metagenome-assembled genomes (MAGs).**

(a) Proportion of variance in MAG-based community composition of sediment communities explained by reef zoning status, sampling trip, GBR sector, and reef site, as inferred by sequential PERMANOVA. (b) The first two components of the MINT sPLS-DA showing clustering by reef zoning status (no-take marine reserves, NTMRs, versus fished reefs) based on indicator MAGs of sediment communities. Points are shaped by GBR sector; ellipses indicate 95% confidence regions. Permutation tests of reef zoning status classification using MINT sPLS-DA based on MAG-based (c) and mOTU-based (d) profiles of sediment communities. Histograms show the null distribution of classification accuracy under leave-one-sector-out cross-validation. Red solid lines indicate the observed classification accuracy; blue dashed lines indicate the 2.5th and 97.5th percentiles of the null distribution.

**Supplementary Tables S1 through S22**

See separate .xlsx file.
